# ARF5/MONOPTEROS directly regulates miR390 expression in the *Arabidopsis thaliana* primary root meristem

**DOI:** 10.1101/463943

**Authors:** Mouli Ghosh Dastidar, Andrea Scarpa, Ira Mägele, Paola Ruiz-Duarte, Patrick von Born, Lotte Bald, Virginie Jouannet, Alexis Maizel

**Affiliations:** Center for Organismal Studies (COS), University of Heidelberg, Im Neuenheimer Feld 230, 69120 Heidelberg, Germany

## Abstract

The root meristem is organized around a quiescent centre surrounded by stem cells that generate all cell types of the root. In the transit amplifying compartment progeny of stem cells further divide prior to differentiation. Auxin controls the size of this transit-amplifying compartment via Auxin Response Factors (ARF) that interact with Auxin Response Elements (AuxRE) in the promoter of their targets. The microRNA miR390 regulates abundance of ARF2, ARF3 and ARF4 by triggering the production of trans-acting (ta)-siRNA from *TAS3*. This miR390/TAS3/ARF regulatory module confers sensitivity and robustness to auxin responses in diverse developmental contexts. Here, we show that miR390 is expressed in the transit-amplifying compartment of the root meristem where it modulates response to auxin. A single AuxRE bound by ARF5/MONOPTEROS (MP) in miR390 promoter is necessary for miR390 expression in this compartment. We show that interfering with *ARF5*/*MP* dependent auxin signaling attenuates miR390 expression in the transit-amplifying compartment. Our results show that ARF5/MP regulates directly the expression of miR390 in the basal root meristem. We propose that ARF5, miR390 and the ta-siRNAs-regulated ARFs are necessary to maintain the size of the transit-amplifying region of the meristem.

**One sentence summary:** The expression of miR390 in the Arabidopsis basal root meristem is controlled by ARF5/MONOPTEROS.

## INTRODUCTION

Growth of the *Arabidopsis thaliana* root is supported by its apical meristem. This root meristem consists of three main regions. The quiescent center (QC), formed by few cells that barely divide, is surrounded by stem cells which divide to form the different cell types that comprise the stereotypical Arabidopsis root. The proximal meristem, located shootward from the QC, is the region where stem cell progeny undergoes rapid, transient-amplifying cell divisions that provide the necessary number of cells for organ growth (Scheres, 2007). Cells leaving the basal meristem undergo rapid cell elongation without division in elongation/differentiation zone (Heidstra and Sabatini, 2014). The size of the meristem is homeostatically regulated by matching the rates of cell production in the meristem and differentiation in the elongation/differentiation zone (Heidstra and Sabatini, 2014). This balance results from the interplay between auxin and cytokinin signaling (Dello Ioio et al., 2007, 2008). The transcriptional auxin signaling relies on Aux/IAA proteins that bind and inhibit Auxin Response Factors (ARFs), DNA-binding proteins in charge of regulating auxin-dependent genes. ARFs bind DNA via an auxin response element (AuxRE) (Boer et al., 2014). In presence of auxin, a complex is formed between Aux/IAA and TIR1/AFB triggering the poly-ubiquitination and subsequent degradation of Aux/IAA, unlocking the ARFs (Paque and Weijers, 2016).

ARF5/MONOPTEROS (MP) plays an essential role in relaying the effects of auxin in multiple developmental contexts (Aida et al., 2002; Bhatia et al., 2016; Hardtke and Berleth, 1998; Przemeck et al., 1996; Smet et al., 2010). MP is essential for the formation of the embryo axis by specifying the root and vasculature. MP is expressed in the lower third of the early embryo and MP loss-of-function prevents the formation of the embryonic root (Weijers et al., 2006). During post-embryonic development, 14 out of the 23 ARFs present in Arabidopsis (Okushima et al., 2005) have been reported to be expressed in the primary root tip (Marin et al., 2010; Rademacher et al., 2011), including MP. Whereas full loss of function *mp* alleles lead to rootless embryos, in weak *mp* allele (*mp-S319*) a root is formed but its growth is impaired (Cole et al., 2009). *ARF3* (Marin et al., 2010), *ARF2*, *ARF8*, *ARF10*, *ARF16, ARF17* (Rademacher et al., 2011), are expressed in the root meristem and their abundance controlled by endogenous small regulatory (s)RNAs (Kasschau et al., 2003; Poethig et al., 2006).

Micro (mi)RNA and trans-acting small interfering RNA (ta-siRNAs) are endogenous small regulatory RNAs that regulate post-transcriptionally the abundance of their target (Bologna and Voinnet, 2014) and control many processes in plants (Mallory and Vaucheret, 2006), in particular root development (Stauffer and Maizel, 2014; Kasschau et al., 2003; Rodriguez et al., 2015; Marin et al., 2010; Yoon et al., 2014, 2010; Carlsbecker et al., 2010; Yu et al., 2015). Of special interest is the *TAS3* ta-siRNA pathway, in which miR390 triggers the biogenesis of ta-siRNAs by ARGONAUTE (AGO)7-mediated cleavage of the non-coding RNA *TAS3*. Subsequent recruitment of SUPPRESSOR OF GENE SILENCING 3 (SGS3) and RNA-DEPENDENT RNA POLYMERASE 6 (RDR6) to the cleaved *TAS3*, results in the DICER-LIKE 4 (DCL4)-dependent production of 21-nt secondary siRNAs called ta-siRNAs targeting members of the *ARF* family (Bologna and Voinnet, 2014). The *TAS3* tasiRNA pathway is conserved across land plants and has been repeatedly employed to regulate ARFs abundance and confers sensitivity and robustness onto the auxin response (Plavskin et al., 2016). In Arabidopsis, tasiRNAs inhibit *ARF2*, *ARF3*, and *ARF4* and functions in the adaxial-abaxial (top-bottom) leaf polarity (Garcia et al., 2006; Adenot et al., 2006; Fahlgren et al., 2006; Hunter et al., 2006), heteroblasty (Hunter et al., 2003), biotic stress response (Cabrera et al., 2016) and lateral root outgrowth (Marin et al., 2010; Yoon et al., 2010). During lateral root formation, the *TAS3* pathway defines an autoregulatory network in which positive and negative feedback regulation of miR390 by ARF2, ARF3, and ARF4 ensures the proper expression of miR390 and maintains *ARFs* concentration in a range optimal for specifying the timing of lateral root growth (Marin et al., 2010; Yoon et al., 2010). The *TAS3* pathway appears thus to be integral to the control of auxin mediated lateral root growth. Interestingly, all components of the pathway are also expressed in the primary root meristem and miR390 has been shown to respond to auxin (Marin et al., 2010; Yoon et al., 2010). In addition miR390 has been suggested to play a role in the auxin induced reduction of the meristem activity (Thimann, 1937; Eliasson et al., 1989; Mähönen et al., 2014; Yoon et al., 2014). Yet, how auxin modulates the expression of miR390 is still unknown.

Here, we show that in the proximal meristem, cells of the transient amplifying compartment express miR390 and that auxin induced reduction of the proximal meristem impacts miR390 expression in the primary root. Ectopic expression of miR390 results in auxin hypersensitivity whereas loss of function results in hyposensitivity. We identify a short segment in the *MIR390a* promoter responsible for the expression of miR390 in the transient amplifying compartment. We show that ARF5/MP interacts directly with this segment via an AuxRE and controls miR390 expression in the proximal meristem. Our data show that miR390 plays a role in the auxin-mediated control of the meristem size and that ARF5/MP is an essential regulator of its expression in the root meristem.

## RESULTS

### MIR390a is expressed in the root meristem where it modulates response to auxin

A transcriptional reporter fusing 2.6kb of sequence upstream of the *MIR390a* (At2g38325) stem loop to GUS shows expression in the primary meristem and the lateral root primordia (Marin et al., 2010). The GUS pattern is identical to the one obtained by whole mount in situ hybridisation (WMISH) for miR390 (Figure 1A, B and (Dastidar et al., 2016)), in agreement with previous results showing that *MIR390a* is the only precursor expressed in the root (Dastidar et al., 2016; Marin et al., 2010). In the primary root meristem, we observe signal in the stem cell niche including the quiescent center, the vascular, epidermal and cortex-endodermal stem cells, but not in the columella stem cells. We also observe a graded expression in the transient amplifying compartment up to the transition zone, where the staining disappears. In the lateral root, the signal is marking the flanks of the primordium (Figure 1A’, B’ and (Marin et al., 2010)).

**Figure 1.**
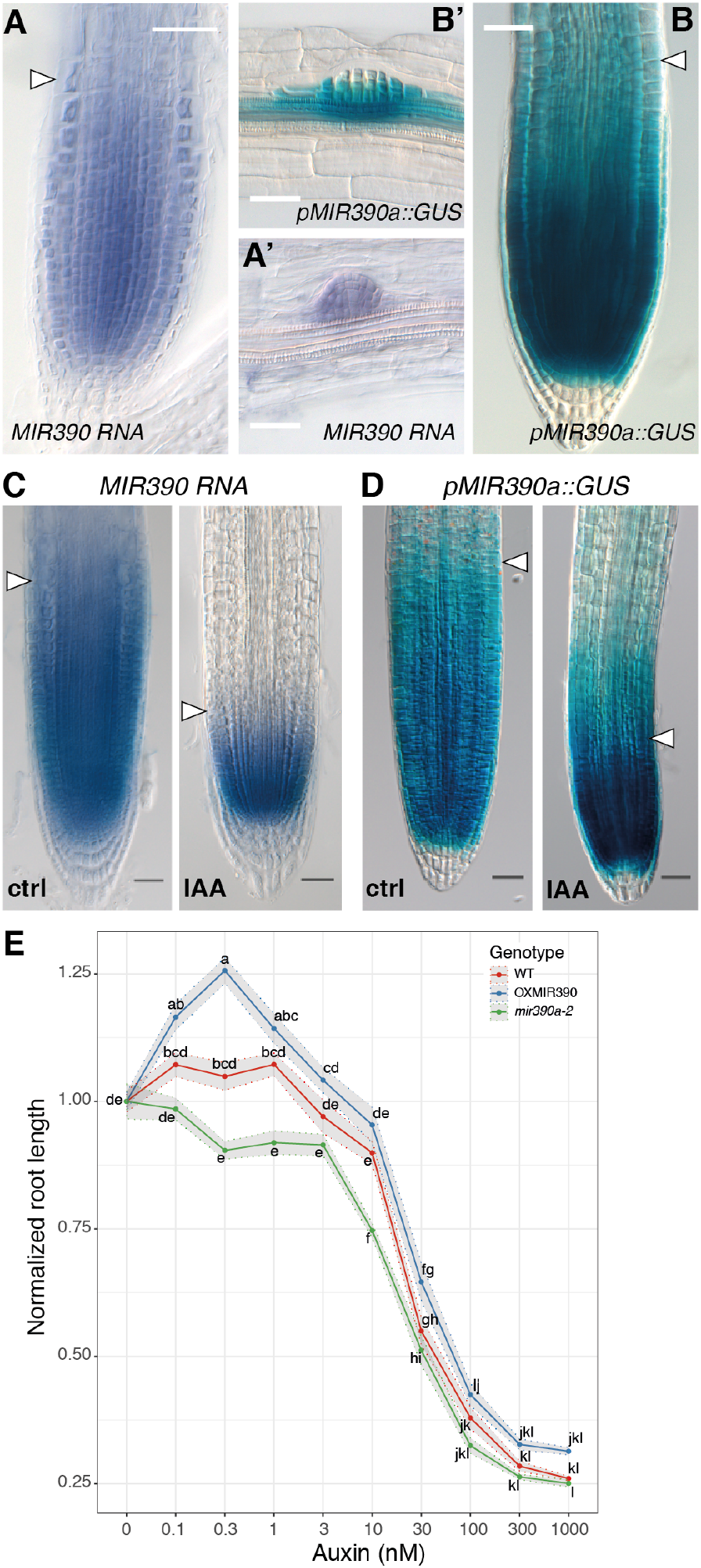
MIR390a is expressed in the root meristem and modulates response to auxin. (A, A’) Expression of miR390 detected by whole mount in situ hybridisation (WMISH) on wild type plants. miR390. (B, B’) expression pattern of the *pMIR390a::GUS* transcriptional reporter for *MIR390a.* miR390 and the MIR390a promoter are expressed in the primary root meristem (A, B) and in lateral root primordia (A’, B’). The arrowhead indicate the transition zone marking the end of the meristem. Images were taken at 5 day after germination (DAG) and scale bars are 25*μ*m. (C, D) Expression of miR390 by WMISH (C) and *pMIR390a::GUS* (D) in response to IAA treatment. Plants (5DAG) were treated by 1*μ*M IAA for 24h before fixation. Images were taken at 6DAG and scale bars are 25*μ*m. (E) Length of the primary root in response to IAA concentrations. Plants were grown for 3 days on control medium before transfer to medium containing the indicating amount of IAA for another 3 days. Root length after 6 days was normalised to its value in absence of exogenous IAA. The lines represent the average root length of 20 plants and the shaded ribbons the standard error to the mean. Comparison between samples was performed using ANOVA and Tukey’s HSD. Samples with identical letters do not significantly differ (α=0.05).

Upon treatment with 1*μ*M auxin (IAA) for 24h, the size of the meristem diminishes by reduction of the transient amplifying compartment (Figure 1C, D) with the concomitant reduction of the expression domain of miR390 (Figure 1C) and its promoter *pMIR390A* (Figure 1D). These data indicate that miR390 is expressed in the meristem, marking the transient amplifying compartment and that auxin and miR390 expression appear to be functionally connected.

The reduction of miR390 expression and accompanying upregulation of the ta-siARF target ARF3 in the primary root meristem has been previously hypothesised to be involved in the inhibition of root growth induced by IAA (Yoon et al., 2014). To test whether miR390 is indeed involved in the regulation of the size of the transient amplifying compartment by auxin, we monitored the IAA inhibition of primary root growth in plants that either constitutively express miR390 (*p35S::MIR390A*, OXMIR390) or with reduced levels of miR390 (*mir390a-2*) (Marin et al., 2010). Plants with lower levels of miR390 (*miR390a-2*, Figure 1E) were more sensitive than wild type to low doses of IAA. Plants with higher levels of miR390 (OXMIR390, Figure 1E) had overall similar response to wild type, albeit with a mild trend toward increased primary root length in low (0.1-1 nM) IAA concentration. This result indicates that miR390 is required for the meristem response to IAA. Based on this result, we asked whether miR390 also controls primary root growth and meristem size in absence of exogenous auxin. We did not observe any difference between *miR390a-2* root length nor meristem size (Supplemental Figure 1). Altogether, these results indicate that miR390 expression in the primary root meristem is modulated by exogenous auxin and that miR390 is involved in the response of the primary root to auxin.

### A 36bp long regulatory element is responsible for the expression of *MIR390a* in the root meristem

Given the role of miR390 in the meristem response to auxin we sought to identify which region of the *MIR390a* promoter is responsible for the expression in the primary root meristem. For this, we generated nested 5’ deletions in the 2.6Kb fragment and fused these to a GUS reporter (Figure 2). For each construct, GUS staining was performed and the presence of signal in the meristem region was scored (n=7 to 20 primary transformants). All reporters containing regulatory sequence up to position −555bp from the transcription start site (+1) had the same expression pattern as the original 2.6Kb long *pMIR390a::GUS* reporter (Figure 2A, B). On the contrary, reporters using 519bp-long or shorter fragments did not present any staining in the meristem (Figure 2A, B). This indicated that the 36bp located between positions −555 and −519 are necessary for expression of the reporter in the primary root meristem. This sequence was dubbed the primary root regulatory element (PRE). Examination of the PRE revealed that it contains a single copy of a putative high affinity auxin response element (CCGACA, AuxRE) (Boer et al., 2014). To test the functional importance of this putative AuxRE to the activity of the PRE and to the expression of *MIR390a* in the root meristem, we deleted this motif in the context of the −555bp reporter (−555∆ARE) and scored for the presence of GUS staining in the meristem of primary transformants. Whereas reporters using the control −555 reporter showed staining of the meristem (77%, n=9), no expression was detected in the meristem of plants transformed with the −555∆ARE reporter (75%, n=12). Taken together, these data indicate that expression of *MIR390a* in the primary root meristem is dependent on a single primary root regulatory element and on the presence of a putative auxin response element therein.

**Figure 2.**
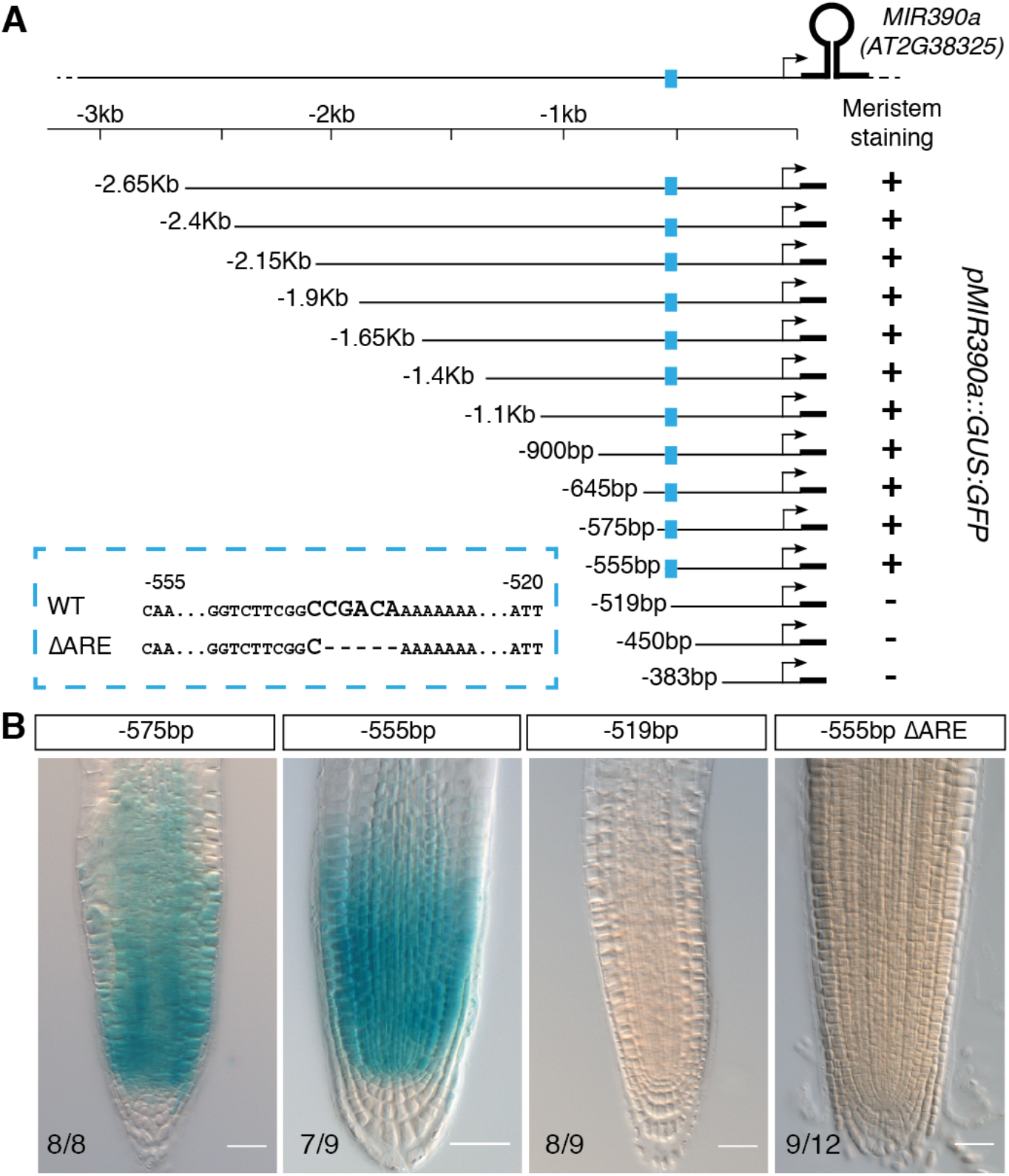
An auxin response element containing enhancer controls MIR390a expression in the root meristem. (A) Nested deletions in the region located upstream of the MIR390a stem loop region. The segment used to drive the expression of the GUS:GFP reporter is represented by a line, the MIR390a transcription start by an arrow and the stem loop by a thick line. The position of the primary root enhancer is indicated in blue and its sequence as well as the putative auxin response element (ARE) are shown in the dashed blue box. (B) Meristem expression of GUS reporter driven by the indicated segment of MIR390a promoter. The numbers indicate the proportion of independent primary transformants displaying the phenotype and scale bars are 25*μ*m.

### A set of five ARFs interact with the primary root regulatory element *ex planta*

The presence of a putative AuxRE in the PRE of *pMIR390a* lead us to seek which ARF could interact with this CCGACA motif. For this we performed a targeted enhanced yeast one hybrid (Y1H) screening (Gaudinier et al., 2011) using a trimerized version of the PRE as a bait and a collection of 16 ARFs as preys. Yeasts expressing ARF4, 5, 8, 9 and 18 grew on selecting medium lacking tryptophan and histidine in presence of 10mM 3AT, whereas growth was much reduced or undetectable in yeasts expressing any of the other ARFs or no ARF at all (Figure 3A). This result indicates that ARF4, 5, 8, 9 and 18 interact with the trimerized PRE. To test whether the interaction required the presence of the AuxRE motif, we repeated the yeast assay expressing this time as a bait a trimerized version of the PRE in which the putative AuxRE has been deleted (PRE∆ARE). In these conditions, yeast expressing ARF4, 5, 8, 9 and 18 were not able to grow (Figure 3B) indicating that interaction between these ARFs and the PRE requires the presence of the ARE.

**Figure 3.**
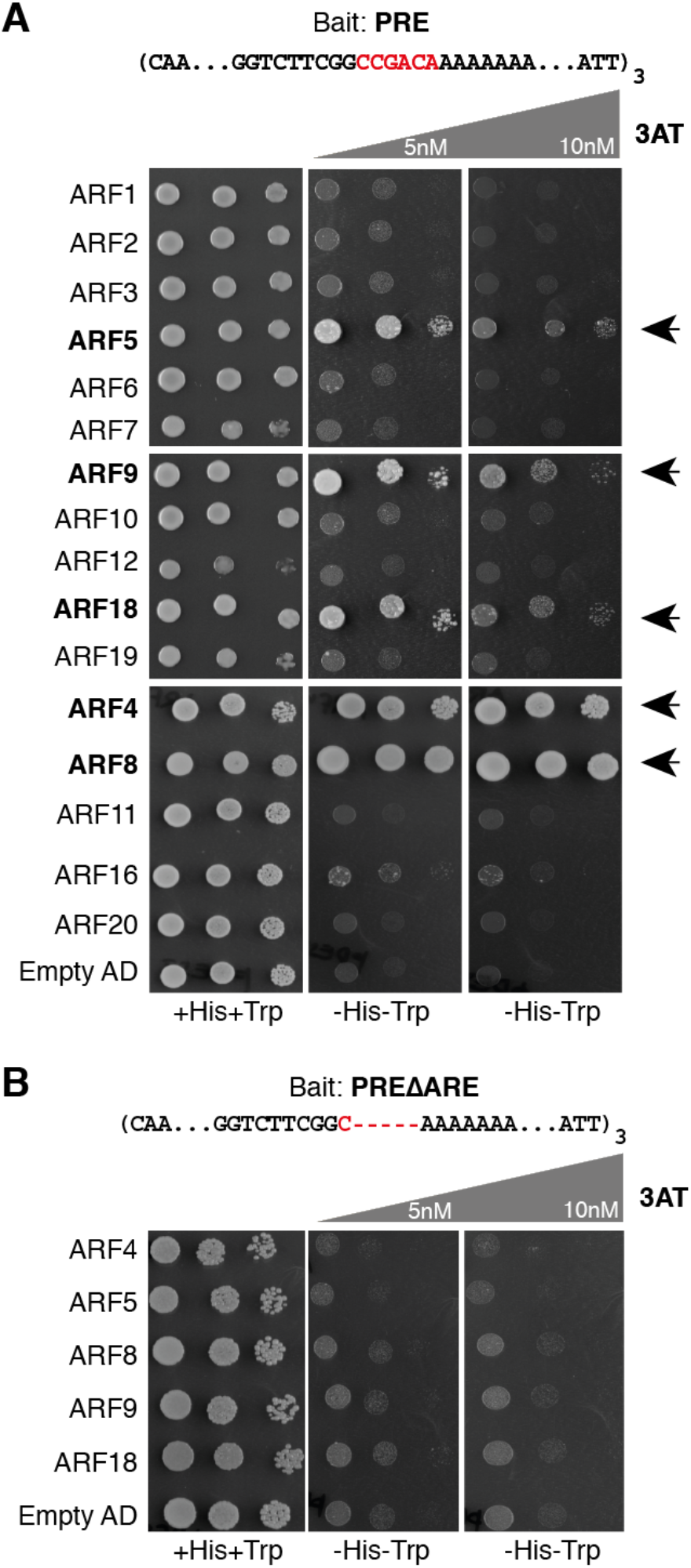
Five ARFs interact with MIR390a primary root enhancer in yeast. (A) Yeast co-expressing a trimerized version of the primary root enhancer (PRE) as bait and a fusion between the indicated ARF and the Gal4 activation domain (AD) were grown on selective media in presence or not of 3AT. ARF5, ARF9, ARF18, ARF4 or ARF8 interacted with PRE resulting in yeast growth on selective medium (arrowheads). (B) Yeast expressing as bait a trimerized version of the PRE lacking the auxin response element (ARE) did not grow on selective medium.

### ARF5/MP controls the expression of miR390 in the root meristem

To identify which of these five ARFs is a bona fide interactor of *pMIR390a* PRE *in planta*, we first examined which of them are expressed in the root meristem. For this, we used previously generated transcriptional reporters (for ARF4 and ARF8) and translational fusions (for ARF5, ARF9 and ARF18) (Rademacher et al., 2011; Marin et al., 2010). The expression pattern of the reporters showed that ARF4, ARF5, ARF8, ARF9 and ARF18 are expressed in the root meristem (Supplemental Figure S2) in domains overlapping with the one of *MIR390a* and could regulate its expression. We then tested if the expression of miR390 is dependent on these ARFs. We quantified the abundance of miR390 by northern blot in plant mutants for the candidate interacting ARFs ARF4, ARF8 and ARF9 as well as ARF2 which did not show interaction with the PRE in yeast (Gutierrez et al., 2009; Marin et al., 2010; Hunter et al., 2006; Tian et al., 2004; Okushima et al., 2005). The levels of miR390 were unaltered between wild type and *arf2*, *arf4*, *arf8* or *arf9* mutants (Fig. 4A). Since strong alleles of *ARF5/MP* lead to defects in axis formation and patterning in embryos and root less seedlings (Hardtke and Berleth, 1998; Donner et al., 2009; Okushima et al., 2005; Schlereth et al., 2010), we crossed the *pMIR390a::GUS* reporter line in the weak *arf5* allele *mpS319* (Cole et al., 2009) and performed GUS staining in the F2 progeny of the cross. Whereas GUS signal was observed at the root tip region of plants heterozygous for *mpS319*, no staining was detected in *mpS319* homozygous plants (Fig. 4B, C). We also performed WMISH and RT-qPCR for miR390 in heterozygous and homozygous *mpS319* seedlings. In both cases, levels of miR390 where strongly reduced in homozygous *mpS319* plants compared to heterozygous (Supplemental Figure S3A, B). Together these results suggest that ARF5/MP is required for miR390 expression in the root meristem.

**Figure 4.**
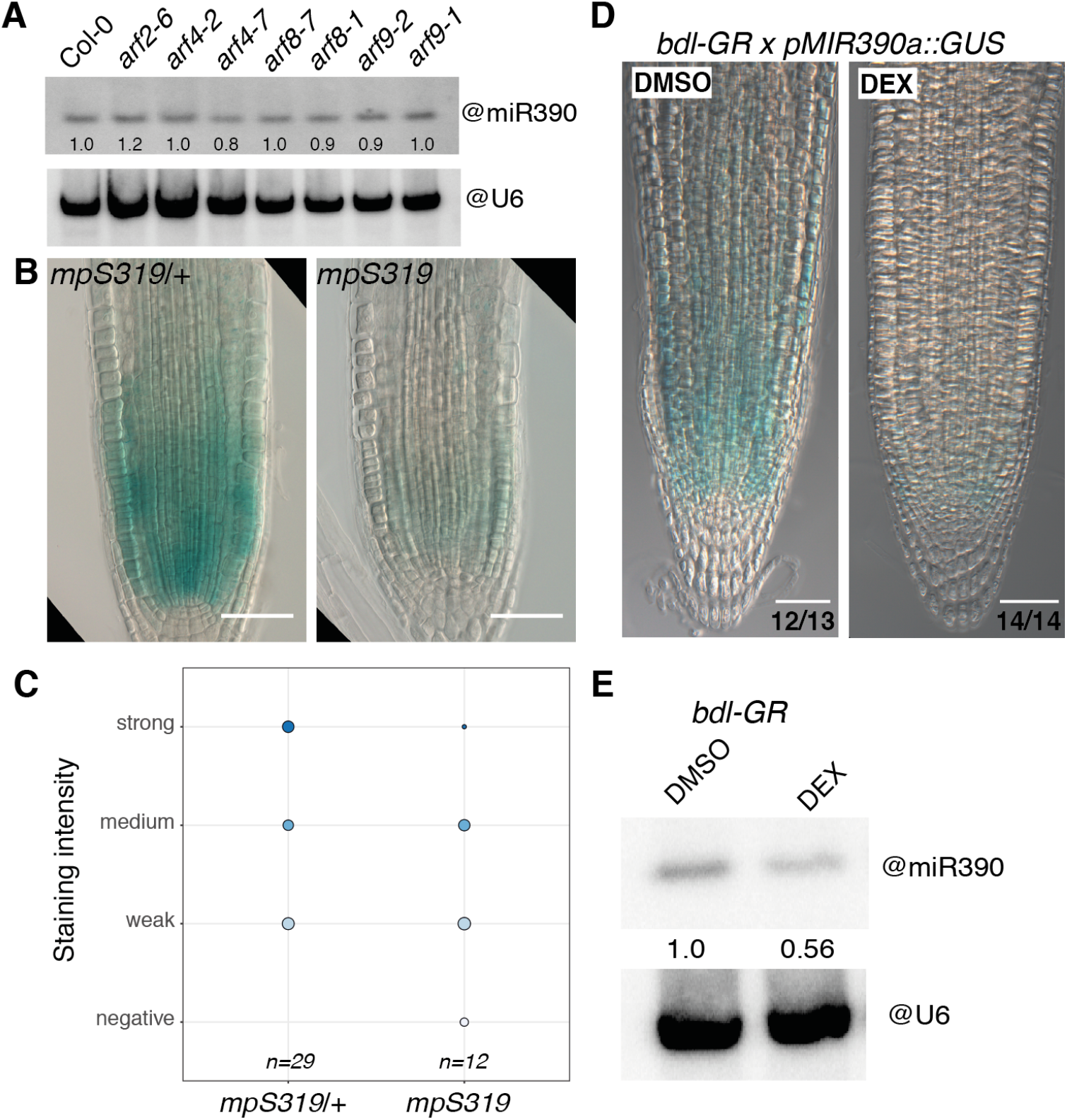
ARF5/MP controls the abundance of miR390 in the primary root meristem. (A) RNA gel blot analysis of 15 *μ*g of root RNA from 7DAG wild-type (Col) or the indicated *ARF* mutant plants hybridized with miR390. U6 snRNA served as a loading control, and numbers are the ratios of miR390 to U6 signal. (B) Expression of *pMIR390a::GUS* in homozygous (*mpS319*) and heterozygous (*mpS319/+*) *monopterous* mutants. Scale bars are 50*μ*m. (C) Distribution of the *pMIR390a::GUS* intensity in homozygous (*mpS319*) and heterozygous (*mpS319/+*)*monopterous* mutants. (D) Expression of *pMIR390a::GUS* in F1 of a cross between *pRBCS5A::bdl-GR* and *pMIR390a::GUS* upon treatment with 10*μ*M dexamethasone (DEX) for 48h to induce the nuclear translocation of *bdl* and inhibition of MP activity or in control conditions (DMSO). The proportion of plants with the depicted signal is indicated. Scale bars are 50*μ*m. (E) RNA gel blot analysis of 15 *μ*g of root RNA from 7-d-old *pRBCS5A::bdl-GR* treated for 48h with 10*μ*M DEX or DMSO and hybridized with miR390. U6 snRNA served as a loading control, and numbers are the ratios of miR390 to U6 signal.

To further test that ARF5/MP is necessary for miR390 expression, we monitored miR390 expression in the *pRBCS5A::bdl-GR* background (Schlereth et al., 2010). In this line a stabilised (and therefore dominant) version of the AUX/IAA BODENLOS/IAA12 which interacts with ARF5/MP is expressed as a GR fusion in the meristem. In absence of dexamethasone (DEX), ARF5/MP activity is normal and consequently meristem organisation and root development are unaltered. Upon DEX treatment, the *bdl* version of IAA12 translocates to the nucleus and inhibits ARF5/MP mediated transcription. We first monitored the expression of the *pMIR390a::GUS* reporter in this inducible knockdown of ARF5/MP. 48 hours after DEX treatment, expression of the *pMIR390a::GUS* reporter was reduced in the root meristem without gross alteration of the organisation of the meristem supporting that ARF5/MP controls expression of the reporter (Figure 4D). We then asked whether the endogenous expression of miR390 was also altered when ARF5/ MP is knocked down. We detected mature miR390 by *in situ* hybridisation (Supplemental Figure S3C) and by northern blot (Figure 4E) in *pRBCS5A::bdl-GR* plants treated or not treated by DEX for 48 hours. We observed a reduction of *in situ* signal at the root meristem (Figure 4E) and a 44% reduction of miR390 signal by northern blot upon DEX treatment (Figure 4F), confirmed by RT-qPCR for miR390 (Supplemental Figure 3D). Together these data indicate that ARF5/MP is required for miR390 expression at the meristem.

### ARF5/MP modulates transcription of MIR390a via the AuxRE in the primary root regulatory element in plant cells

We found that ARF5/MP interacts in yeast with *pMIR390a* via the auxin response element located in the regulatory element required for expression of miR390 in the root meristem and that, genetically, ARF5/MP is required for miR390 expression in this region. To test whether ARF5/MP is able to directly induce the *MIR390a* promoter in plant cells, we co-expressed in tobacco leaves ARF5/MP with different fragments of the *MIR390a* promoter. To circumvent the need to trigger auxin-mediated degradation of AUX/IAA to activate ARF5/MP, ARF5/MP was expressed as a fusion with the VP16 activation domain. Co-expression of ARF5/MP-VP16 with the 555bp long fragment of *MIR390a* promoter fused to LUC was sufficient for expression (Figure 5A). When ARF4-VP16 or no ARF were co-expressed, reduced (ARF4) or no significant expression of LUC could be detected (Figure 5A). Variants of the promoter in which the PRE is not present (519bp and 94bp long segments) or elimination of the AuxRE element in the 555bp long one (−555∆ARE) reduced strongly the levels of LUC expression in presence of ARF5/MP-VP16 (Figure 5A). Together, these results indicate that ARF5/MP is able to stimulate transcription from the *MIR390a* as long as it contains the PRE and its AuxRE.

**Figure 5.**
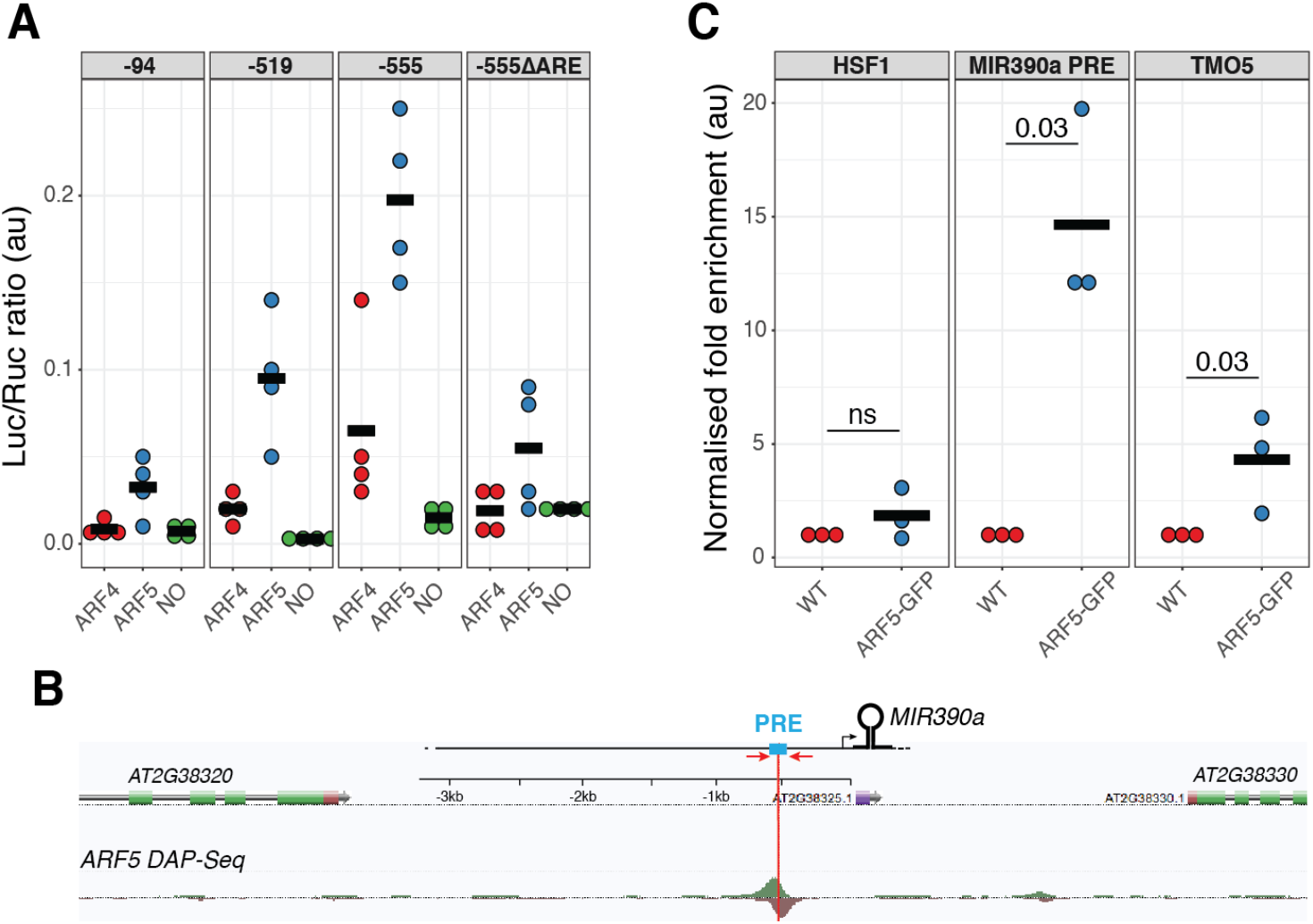
ARF5/MP activates the *MIR390a* promoter via the auxin response element located in the primary root enhancer. (A) Fusions between the VP16 domain and ARF4 (ARF4), ARF5 (ARF5) or empty plasmid (NO) were co-expressed in tobacco leaves with different segment of the MIR390a promoter (−94bp, −519bp or −555bp see Figure 2) driving the expression of the firefly luciferase (Luc). In the −555∆ARE promoter, a 5bp deletion removes the ARE in the −555bp segment (Figure 2). A 35S-driven renilla luciferase (Ruc) is used to normalise activity (Luc/Ruc, expressed in arbitrary units). Each dot is a biological replicate and the horizontal bars the mean of the four replicates. (B) Annotated screen shot of genome browser at the MIR390a locus with a ARF5 DAP-seq peak {OMalley:2016kh} located at the same position as the PRE (blue box). Position of the primers used for ChIP-qPCR (C) is depicted by red arrows. (C) Enrichment of ARF5-GFP at three loci was assessed by ChIP-qPCR. Chromatin prepared from wild type (WT) or ARF5-GFP plants was immunoprecipitated by an anti-GFP antibody and enrichment of ARF5-GFP in the region PRE of *MIR390a* (MIR390a PRE), in the second exon of TMO5 or upstream of the HSF1 loci was assessed by qPCR. Fold enrichment in the immunoprecipitated fraction over input fraction in ARF5-GFP sample was normalised to the one in WT plants and expressed as arbitrary unit. Each dot is a biological replicate and the horizontal bars the mean of the three replicates. Statistical significance was evaluated by the Kruskal-Wallis test and p values are indicated.

### ARF5/MP interacts with the primary root regulatory element of *MIR390a* in Arabidopsis

Mining of the Arabidopsis cistrome data (O’Malley et al., 2016)e revealed the presence of a DAP-seq peak for ARF5/MP ~550bp upstream of *MIR390a* transcription start which corresponds exactly to the location of the PRE regulatory element we identified (Figure 5B and 2A). This further substantiates that ARF5/MP interaction with this regulatory element might be relevant. To validate that ARF5/MP interacts with *MIR390a* in Arabidopsis, we performed a chromatin immunoprecipitation assay (ChIP) using a functional ARF5:GFP fusion protein driven by the native ARF5 promoter (ARF5-GFP) (Schlereth et al., 2010). Control immunoprecipitations were performed on Col-0 (WT). Promoter fragments encompassing the PRE region were found to be ~15 fold enriched in ARF5-GFP over WT (Figure 5C, p=0.03, Kruskal Wallis with n=3 biological replicates). Under the same conditions, ARF5-GFP was enriched ~5 fold at the *TARGET OF MONOPTEROS5* (*TMO5*) locus (Schlereth et al., 2010), which served as a positive control. No enrichment was observed at the unrelated locus Heat shock factor 1 (HSF1) (Figure 5C and Supplemental Figure S3). This result indicates that ARF5/MP interacts with *MIR390a* promoter in the PRE region.

## DISCUSSION

Here, we show is expressed in the transit amplifying compartment responsible for the radial patterning of the root and its growth. This expression is controlled by a single enhancer and the high affinity auxin response element (AuxRE) it contains. We show that ARF5/MP interacts with this enhancer via the AuxRE in yeast and in plant cells and that in planta ARF5/MP binding is enriched in the region of MIR390a promoter containing the enhancer. By interfering with ARF5/MP dependent auxin signaling we show that expression of the miR390 is reduced, showing that ARF5/ MP is necessary for miR390 expression in this compartment of the meristem.

Like their protein-coding gene counterparts, transcriptional regulation of miRNA gene involves regulation by cis-regulatory elements and trans regulators. Although hundreds of miRNA have been identified in plants, there is only a handful of studies linking expression of miRNA to specific transcription factors. An in silico approach has shown that cis-regulatory elements corresponding to binding motifs for transcription factors such as LEAFY (LFY), ARFs, and AtMYC2 are overrepresented in the putative promoter region of MIRNA genes (Megraw et al., 2006; Megraw and Hatzigeorgiou, 2010) but the functional relevance of these enrichment has not been tested. Timing of phase transition and flowering relies on the sequential activation of the miR156/SPL and miR172/AP2 regulatory networks (Wu et al., 2009; Wang et al., 2009). The miR156-regulated SPL9 and SPL10 transcription factors promote the expression of the miR156 in a feed-forward loop as well as the sequential expression of miR172 to promote floral transition (Wu et al., 2009; Wang et al., 2009). Transcription of miR398 in Arabidopsis is induced in response to copper starvation and is involved in the degradation of mRNAs encoding copper/zinc superoxide dismutase (Sunkar et al., 2006). The transcription factor SPL7 is required for the expression of miR398 and other copper-responsive miRNA (miR397, miR408, and miR857). SPL7 binds directly to GTAC motifs in the miR398 promoter in vitro, and these motifs were essential and sufficient for the response to copper deficiency in vivo (Yamasaki et al., 2009). Our results indicate that miR390 is one of ARF5/MP target in the meristem. miR390 is the trigger of a regulatory network fine tuning the abundance of ARF2, ARF3, ARF4 and therefore ensuring sensitivity and robustness to auxin signaling in several developmental contexts (Garcia et al., 2006; Adenot et al., 2006; Marin et al., 2010; Fahlgren et al., 2006; Hunter et al., 2003, 2003; Cabrera et al., 2016; Marin et al., 2010; Yoon et al., 2010). This regulatory network is evolutionary conserved (Plavskin et al., 2016) and characterized by a retrocontrol of the ta-siRNA-controlled ARF on the expression of the miR390 (Plavskin et al., 2016; Marin et al., 2010). We identified ARF5/MP as an upstream regulator of miR390 on the root meristem. Whereas ARF5/MP is not a target of the miR390/TAS3 module, it is interesting that an ARF controls the expression of the miR390, suggesting that the miR390/TAS3/ ARF regulatory network is an integral part of the regulatory networks mediating auxin response. It would be interesting to study whether in other developmental contexts where the miR390/TAS3/ARF regulatory network has been implicated such as lateral root and leaf patterning, similar network motif involving ARF5/MP or other ARF have also been coopted to regulate miR390 expression. During lateral root formation, ARF3 and ARF4 are respectively repressor and inducer of miR390 (Marin et al., 2010). Whereas we identified ARF4 as interacting with the AuxRE present in the primary root enhancer in yeast, ARF3 was not. It would be interesting to map the cis-regulatory motif(s) controlling the expression of miR390 in the lateral root to see whether an AuxRE is present and which ARF interact with it.

Reduction of miR390 levels leads to hypersensitivity of the meristem to the inhibition of meristem growth induced by mild exogenous concentrations of auxin, whereas increased miR390 levels have opposite effects. miR390 therefore acts as a modulator of the effects of auxin on the size of the transit amplifying compartment. Previous work has hypothesized, based on the modulation of expression patterns, that the miR390-*TAS3*-*ARF3* module could control differentially the behavior of the meristem in response to exogenous auxin (Yoon et al., 2014). Here, we provide experimental evidences that levels of miR390 modulate the responsiveness of the meristem to auxin.

ARF2, ARF3 and ARF4, promote primary root growth (Marin et al., 2010) and are pot-transcriptionally regulated by miR390/TAS3. A mutation in miR390 does not dramatically alter neither the size nor the growth capacity of the meristem in absence of exogenous auxin suggesting that the effects of the miR390/TAS3 module in the primary root may be buffered by additional control mechanisms. This is coherent with the absence of effect of gain and loss of function in *TAS3* on primary root length (Marin et al., 2010). At least one other miRNA-controlled regulatory network has been involved in the control of root meristem size. miR396 is transcribed in the QC and columella and non-cell-autonomously represses a set of GROWTH REGULATING FACTORs (GRFs) which are transcription factors localised exclusively in the transit amplifying compartment where they locally promote cell division. miR396 ensures the exclusion of these GRFs from the stem cell niche and contribute to the transition between the stem cell niche and transit amplifying compartment of the root meristem (Bazin et al., 2013; Rodriguez et al., 2015). The miR396/GRF and miR390/TAS3/ARF3 regulatory networks both control leaf development (Rodriguez et al., 2010; Garcia et al., 2006; Adenot et al., 2006; Fahlgren et al., 2006; Hunter et al., 2006) and have been shown to genetically interact (Mecchia et al., 2013). It would be interesting to investigate if both modules also interact in the root meristem.

## MATERIALS & METHODS

### Plant Materials and Growth Conditions

*Arabidopsis thaliana* accession Col-0 was used throughout this study. For transient assays *Nicotiana benthamia*na were used. The *miR390a-2* mutant (WiscDsLox440F06) was described in a(Marin et al., 2010)m. The *arf8-1,arf8-7,arf9-1,arf9-2, arf4-7* mutants were described in (Hunter et al., 2006; Gutierrez et al., 2009; Marin et al., 2010; Tian et al., 2004; Okushima et al., 2005), *MP-GFP (MP::MP-GFP)*and *pRPS5A::bdl-GR* in (Schlereth et al., 2010), *mpS319* in (Cole et al., 2009), the *pARF4::GFP, pARF8::GFP*, *ARF9:GFP* and *ARF18:GFP* in (Marin et al., 2010; Rademacher et al., 2011). *Arabidopsis thaliana* were grown on soil in controlled plant rooms at 23°C under long day conditions (16h day length, LED illumination 150mmol quanta m^-2^ s^-1^). *Nicotiana benthamiana* were grown on soil at 25°C with 16h day length (150mmol quanta m^-2^ s^-1^). For *in vitro* growth on plates, seeds were surface sterilized with 70% EtOH+0.1% SDS for 10-15 mins followed by washing 3 times with 99% EtOH and sown out on 1/2 MS (Murashige-Skoog) medium plates and 2.3 mM MES (pH 5.8) in 0,8% phytoagar. After stratifying the seeds in the dark (4°C) for 2 to 3 d, plates were placed in a vertical orientation inside growing chambers (23°C, 150mmol quanta m^-2^ s^-1^). For chemical treatments, DEX (D4902-1G; Sigma-Aldrich) was stored as 30 mM stocks in DMSO and used at 10 *μ*M for the indicated periods. For auxin sensitivity assays, plants were initially grown for 3 d before transferred onto fresh plates containing the indicated concentrations of IAA (I3750-25G-A; Sigma-Aldrich).

### Statistical Analysis and Plotting

All statistical analysis and plotting were performed with R (www.r-project.org) and plotting with the ggplot2 package (v3.0.0.900).

### Analysis of Root Growth

For root elongation measurements, seedlings were grown vertically for 10 days. Starting from day 3 after germination until the end of the experiment at day 10, a dot was drawn at the position of the root tip. Finally, plates were scanned on a flatbed scanner, and the root length was measured over time with Fiji (Schindelin et al., 2012). Meristematic zone length was determined according to the file of cortex cells from confocal microscopy images after mPS-PI treatment (Truernit et al., 2008). The meristematic zone was defined as the region of isodiametric cells from the QC up to the cell that was twice the length of the immediately preceding cell (Dello Ioio et al., 2008).

### Cloning and Generation of Transgenic Lines

The p35S::MIR390 construct was obtained by Gateway cloning by PCR amplification of the *MIR390b* stem loop (AT5G58465) cloning in pDONR201 and recombination in pAM506 (p35S::GateWay_RfA:term). Nested deletions in pMIR390a were obtained by Gateway cloning. Each fragment was amplified by PCR, cloned in pJLSmart and recombined in pKGWFS7 (Karimi et al., 2007). Deletion of the AuxRE in the −555 segment was obtained by gene synthesis and cloned as before. For the Y1H, the coding regions of ARF*4,8,11,16,17,20* were amplified from total seedling cDNA cloned in pDONR221 or pCR8 and Gateway cloned in AD-2μ destination vector by a Gateway LR reaction (Gaudinier et al., 2011). Trimerised version of the 36bp PRE bait (with and without ARE) were obtained by gene synthesis and cloned in the yeast 1 hybrid bait vectors, pMW#2 containing the *HIS3* reporter gene and integrated in the strain (YM4271) (Deplancke, 2004). VP16-fusions for ARF4 and ARF5 were generated by GreenGate cloning (Lampropoulos et al., 2013) with the following modules pUB10 (A), VP16 (B), ARFs coding regions (C), HA-tag (D) and UB10term (E) in pGGZ003. Fragments of pMIR390a were Gateway cloned upstream of LUC in pLUC_GW vector. All primers are listed in Supplemental Table 1.

### GUS (β-glucuronidase) assay, *in situ* hybridization and Microscopy

GUS activity was carried out at 37°C overnight (16h) using 2 mM ferri/ ferrocyanide as described (Weigel and Glazebrook, 2002). After GUS staining whole seedlings were cleared and mounted on 50% glycerol and detected by light microscopy using Differential Interference Contrast (DIC) on a Zeiss Axio Imager M1 (Carl Zeiss, Göttingen, Germany) microscope using a Plan-Apochromat 20X/1.4 NA objective. Whole mount *in situ* hybridizations for miR390 were performed exactly as described (Dastidar et al., 2016). Confocal laser scanning microscopy was performed throughout the study using a Plan Apochromat 20x, 0.8-NA lens on a Leica SP8 or SPE microscopes. Roots were stained with 10 *μ*g/mL propidium iodide for 2 min, rinsed, mounted in water, and visualized after excitation by an argon 488-nm laser line. The fluorescence emission was collected from 590 to 700nm (propidium iodide), 496 to 542nm (GFP).

### RNA isolation and RNA blot

For RNA isolation phenol-chloroform extraction procedure was carried out as describe(Marin et al., 2010). Small RNA northern blot analysis was performed as described (Marin et al., 2010). RNA gel blots were hybridized with the miR390 probe together with U6 probe as a loading control. Non-saturated signals were quantified on a Fuji FLA 7000 scanner.

### miRNA Expression Analysis via RT-qPCR

Total RNA was isolated from root tissue as described above. Total RNA (2*μ*g) was treated with RNase-free DNase I (Fermentas). First-strand cDNA synthesis was performed using SuperScript II Reverse Transcriptase (Invitrogen). Five times diluted cDNA was used for amplification. A parallel reaction without reverse transcriptase enzyme was used as a control for genomic DNA contamination. Quantitative PCR was performed using a modified protocol to detect mature microRNA (Chen, 2005) with a BioRad thermocycler using SYBR Green I (Roche) to monitor double-stranded DNA synthesis. The relative transcript level was determined for each sample and normalized (Livak and Schmittgen, 2001) using the PROTEIN PHOSPHATASE2A cDNA level (AT1G13320, athRef1) (Czechowski et al., 2005). Melting curve analyses at the end of the process and “no template controls” were performed to ensure product-specific amplification without primer-dimer artefacts. Primer sequences are given in Supplemental Table 1.

### Dual-luciferase reporter assay

Tobacco plants were infiltrated with the indicated combination of VP16-fused ARF and *pMIR390a::LUC* constructs along a *p35S::renillaLuciferase.* 48 hours post infiltration detection of luciferases was performed the Dual-luciferase reporter assay system (Promega) and a TECAN Infinite M1000 plate reader.

### Yeast-one-hybrid assay

The eY1H protocol previously described (Gaudinier et al., 2011) was followed. Coding regions of *ARF 4, 8, 11, 16, 17, 20* were cloned in the lab whereas vector containing *ARF1, 2, 3, 5, 6, 7, 9, 10, 12, 18, 19* were obtained from S. Brady (UC,Davis,USA).

### ChIP

ChIP experiments were performed as described in (Gendrel et al., 2005) with a few modifications. One to two grams of *A. thaliana* root tissue from 7 day-old wild type and MP-GFP plants were used. Fixation and cross-linking by 1% formaldehyde was performed twice for 10 mins under vacuum. Tissue was rinsed thoroughly in water, dried, frozen in liquid nitrogen and ground. The resulting powder was resuspended in 30mL of extraction buffer 1 for 10 minutes at 4°C before being filtered twice on 90μm and 50μm meshes then centrifuged at 3,000g for 20 mins at 4°C. The pellet was resuspended in 300μL extraction buffer 2 and centrifuged at 12,000g for 10 mins at 4°C. The pellet was resuspended in 300μL extraction buffer 3 and centrifuged at 16,000g for 1h at 4°C. The resulting pellet was then lysed with 300μL of nuclei lysis buffer and then sonicated for 10 mins. The debris were pelleted by centrifugation at 12,000g for 5 mins at 4°C. At this point 10μL of the sample was set aside as ‘Input control’ and the rest was used for immunoprecipitation after diluting 10 times with ChIP dilution buffer. The samples were divided in different tubes and incubated for 1hr with gentle agitation with 40 μL of Protein A agarose beads (Invitrogen). After spinning down the beads the pre-cleared supernatant was incubated overnight with 0.5 μL anti-GFP antibody (ChIP Grade; Abcam ab290) and Protein A agarose beads at 4°C with rotation. Beads were centrifuged at 3,800g at 4 °C for 30 seconds and successively washed in low salt wash buffer, high salt wash buffer, LiCl buffer and TE buffer. Immune complexes were eluted from the beads by incubation in 250 μL elution buffer at 65°C for 15 mins with intermittent shaking. After beads were pelleted, proteins were reverse cross-linked by adding 20 μL of 5M NaCl at 65°C overnight. All samples were treated with 2 μL of Proteinase K (10mg/mL) and DNA extracted using Qiagen mini elution kit and dissolved in 40μL of water. Fold enrichment at specific loci was quantified by qPCR on 1μL of each sample with the respective primers, as ratio of the immunoprecipitated fraction over input in MP-GFP normalised to the same ratio in WT. The primers used in the ChIP experiments are listed in Supplemental Table 1.

## ACKNOWLEDGEMENTS

We thank C. Bellini for the *arf9* and *arf8* mutants, J. Lohmann for the *MP-GFP* reporter, D. Wagner for the *pRPS5A::bdl-GR* seeds, S. Brady for the eY1H system. This work was supported by the DFG (MA5293/2), the Land Baden-Württemberg, the Chica und Heinz Schaller Stiftung, the CellNetworks cluster of excellence and the Boehringer Ingelheim Foundation.

## CONFLICT OF INTEREST STATEMENT

The authors declare no conflict of interest.

## SUPPORTING INFORMATION

The following materials are available in the online version of this article.

- Supplemental Figure 1 (related to Figure 1) Root growth and meristem of *mir390a-2*
- Supplemental Figure 2 (related to Figure 3) Expression of the PRE-interacting candidates ARFs in the root meristem
- Supplemental Figure 3 (related to Figure 4). Reduction of miR390 levels upon inhibition of ARF5/MP
- Supplemental Figure 4 (related to Figure 5). ARF5 DAP-Seq at TMO5 and HFS1 loci
- Supplemental Table 1: list of primers used.

## Supplemental Material

**Supplemental Figure 1 (related to Figure 1).**
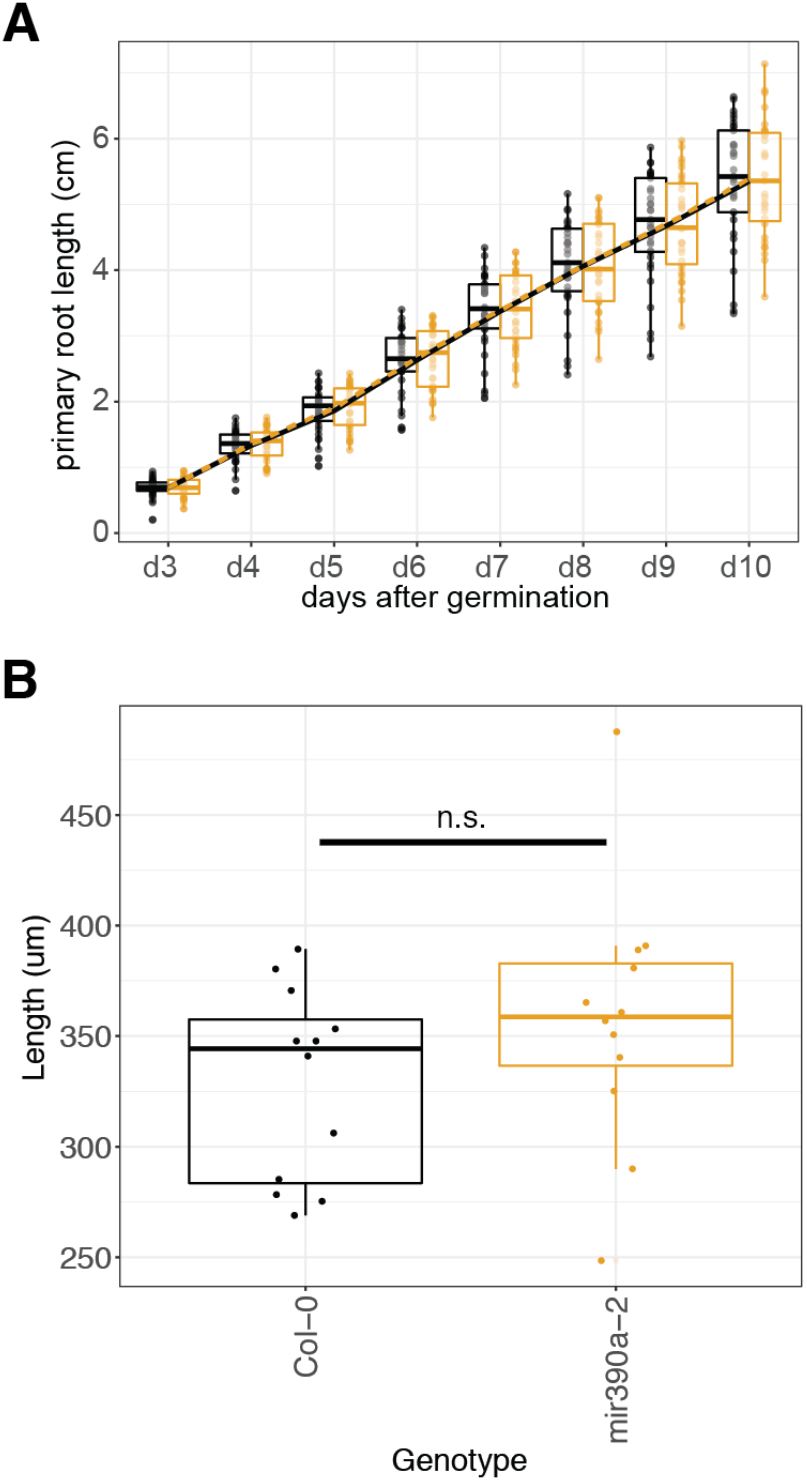
Root growth and meristem of *mir390a-2*. (A) Root elongation (cm) in wild type and *mir390a-2*. Box plots represent the distribution of 32 plants. (B) Meristematic zone length (μm) in wild type and *mir390a-2*. Box plots represent the distribution of 12 plants. In the box plots, the thick line represent the mean of the distribution, the box the 25th-75th interquartile range, the lower whisker the 5th quartile and the upper one the 95th quartile.

**Supplemental Figure 2 (related to Figure 3).**
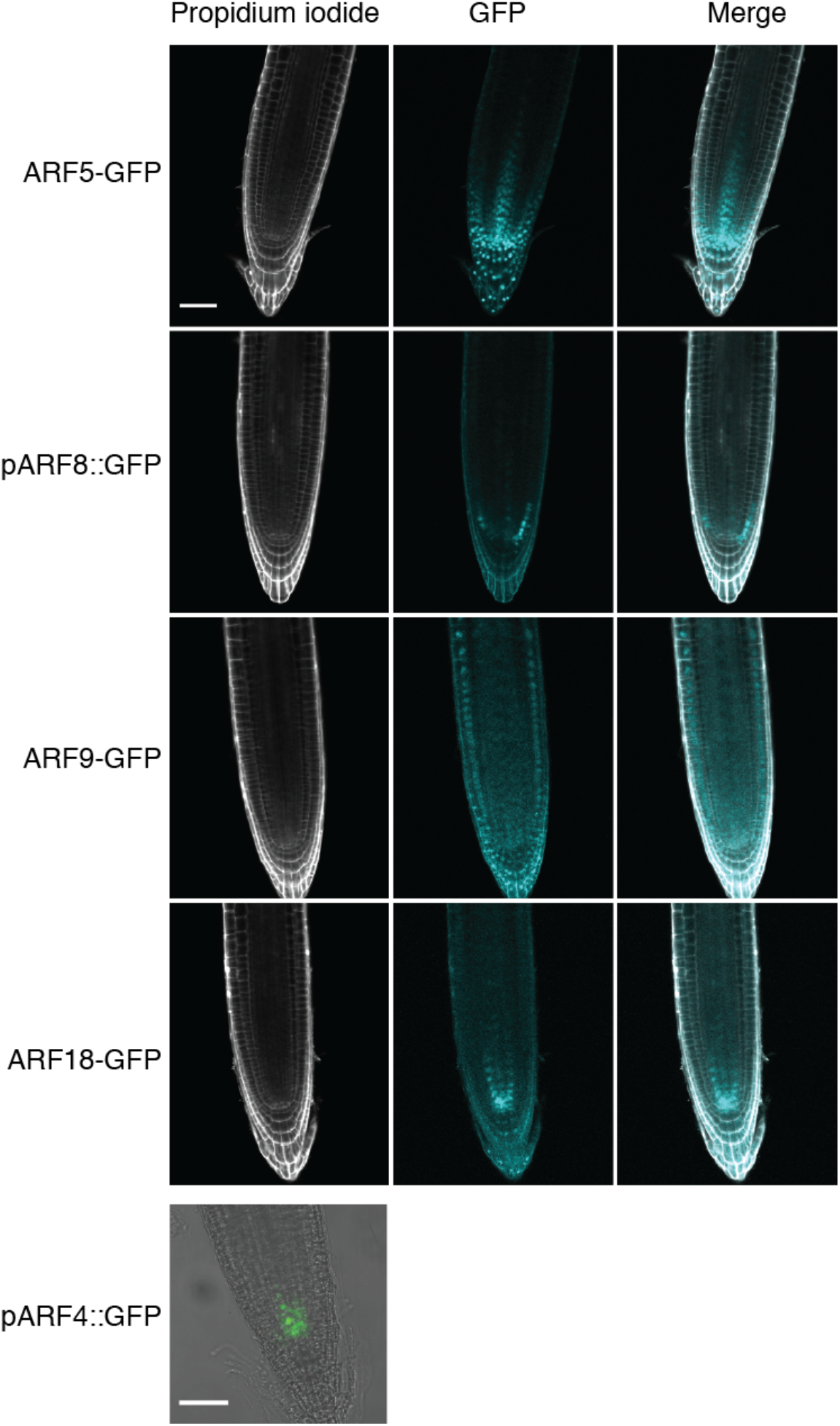
Expression of the PRE-interacting candidates ARFs in the root meristem. Confocal sections of the indicated translational (ARF5-GFP, ARF9-GFP, ARF18-GFP) or transcriptional (pARF8::GFP, pARF4::GFP) fusions.

**Supplemental Figure 3 (related to Figure 4).**
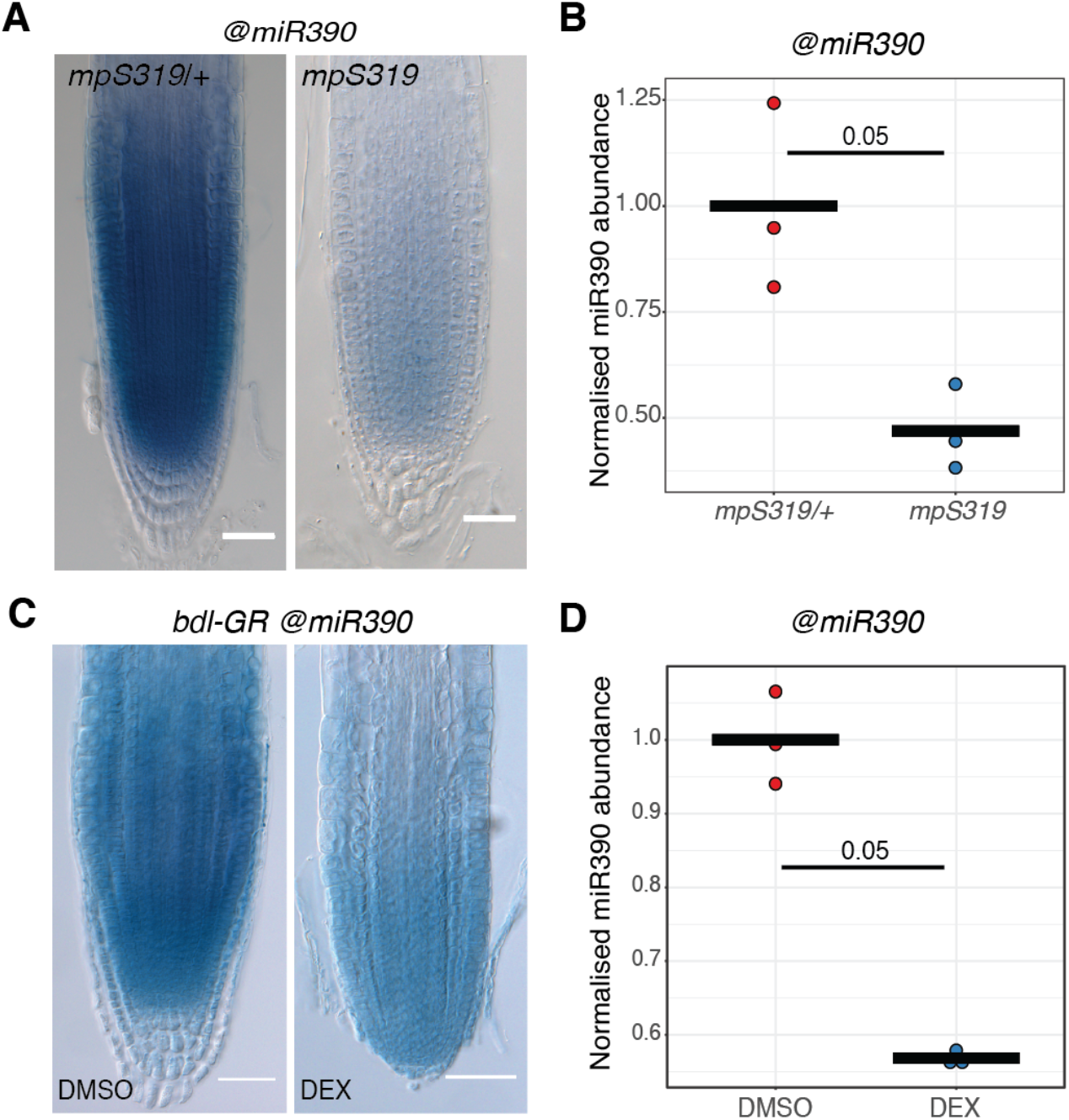
Reduction of miR390 levels upon inhibition of ARF5/MP. (A) Expression of miR390 by WMISH in *mpS319* homozygous and heterozygous (*mpS319/+*). Images were taken at 6DAG and scale bars are 50μm. (B) RT-qPCR analysis of miR390 levels in *mpS319* homozygous and heterozygous (*mpS319/+*) plants. Each dot represent the abundance of miR390 normalised to athRef1 (AT1G13320) in a biological replicate and the horizontal bars the mean of the three replicates. Statistical significance was evaluated by the Kruskal-Wallis test and p values indicated. (C) Expression of miR390 by WMISH in *bdl-GR* plants treated with 10μM DEX or DMSO as control for 3 days. Images were taken at 6DAG and scale bars are 50μm. (D) RT-qPCR analysis of miR390 levels in *bdl-GR* plants treated with 10μM DEX or DMSO as control for 24h. Each dot represent the abundance of miR390 normalised to athRef1 (AT1G13320) in a biological replicate and the horizontal bars the mean of the three replicates. Statistical significance was evaluated by the Kruskal-Wallis test and p values indicated.

**Supplemental Figure 4 (related to Figure 5).**
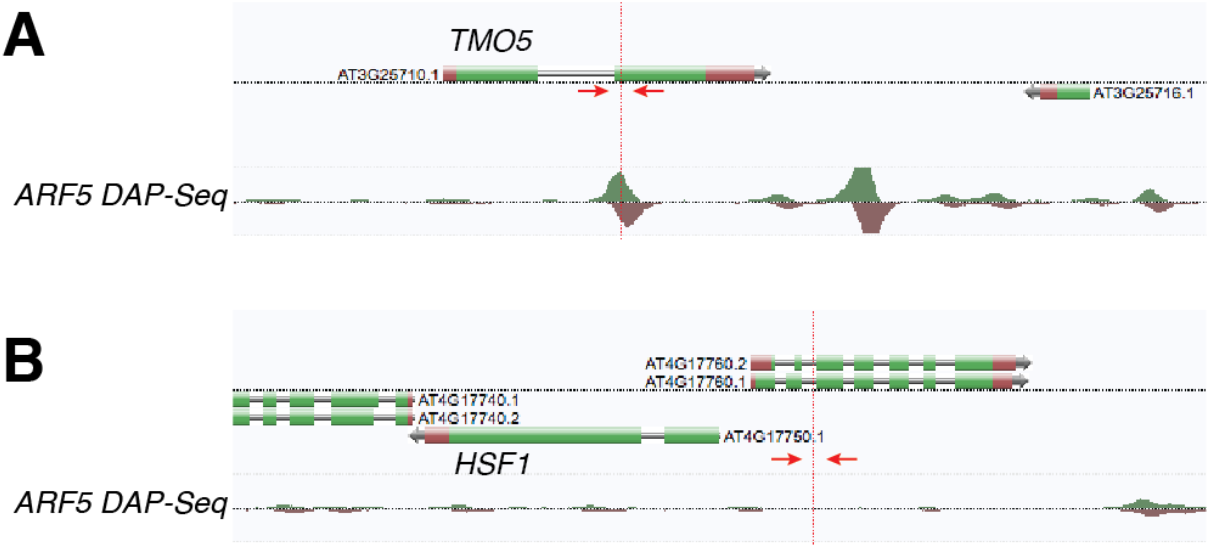
ARF5 DAP-Seq at TMO5 and HFS1 loci. Annotated screen shot of genome browser at the TMO5 (A, AT3G25710) and the HSF1 (B, AT4G17750) locis with a ARF5 DAP-seq peaks (O’Malley et al. 2016). Position of the primers used for ChIP-qPCR (Figure 5C) at each locus is depicted by red arrows.

## BIBLIOGRAPHY

Adenot, X., Elmayan, T., Lauressergues, D., Boutet, S., Bouché, N., Gasciolli, V., and Vaucheret, H. (2006). DRB4-dependent TAS3 trans-acting siRNAs control leaf morphology through AGO7. Curr. Biol. 16: 927–932.

Aida, M., Vernoux, T., Furutani, M., Traas, J., and Tasaka, M. (2002). Roles of PIN-FORMED1 and MONOPTEROS in pattern formation of the apical region of the Arabidopsis embryo. Development 129: 3965–3974.

Bazin, J., Khan, G.A., Combier, J.-P., Bustos-Sanmamed, P., Debernardi, J.M., Rodriguez, R., Sorin, C., Palatnik, J., Hartmann, C., Crespi, M., and Lelandais-Brière, C. (2013). miR396 affects mycorrhization and root meristem activity in the legume Medicago truncatula. The Plant Journal 74: 920–934.

Bhatia, N., Bozorg, B., Larsson, A., Ohno, C., Jönsson, H., and Heisler, M.G. (2016). Auxin Acts through MONOPTEROS to Regulate Plant Cell Polarity and Pattern Phyllotaxis. Curr Biol 26: 3202–3208.

Boer, D.R., Freire-Rios, A., van den Berg, W.A.M., Saaki, T., Manfield, I.W., Kepinski, S., López-Vidrieo, I., Franco-Zorrilla, J.M., de Vries, S.C., Solano, R., Weijers, D., and Coll, M. (2014). Structural basis for DNA binding specificity by the auxin-dependent ARF transcription factors. Cell 156: 577–589.

Bologna, N.G. and Voinnet, O. (2014). The Diversity, Biogenesis, and Activities of Endogenous Silencing Small RNAs in Arabidopsis. Annu Rev Plant Biol 65: 473–503.

Cabrera, J., Barcala, M., García, A., Rio-Machín, A., Medina, C., Jaubert-Possamai, S., Favery, B., Maizel, A., Ruiz-Ferrer, V., Fenoll, C., and Escobar, C. (2016). Differentially expressed small RNAs in Arabidopsis galls formed by Meloidogyne javanica: a functional role for miR390 and its TAS3-derived tasiRNAs. New Phytologist 209: 1625–1640.

Carlsbecker, A. et al. (2010). Cell signalling by microRNA165/6 directs gene dose-dependent root cell fate. Nature 465: 316–321.

Chen, C. (2005). Real-time quantification of microRNAs by stem-loop RT-PCR. Nucleic Acids Research 33: e179–e179.

Cole, M., Chandler, J., Weijers, D., Jacobs, B., Comelli, P., and Werr, W. (2009). DORNROSCHEN is a direct target of the auxin response factor MONOPTEROS in the Arabidopsis embryo. Development 136: 1643–1651.

Czechowski, T., Stitt, M., Altmann, T., Udvardi, M.K., and Scheible, W.-R. (2005). Genome-wide identification and testing of superior reference genes for transcript normalization in Arabidopsis. Plant Physiol. 139: 5–17.

Dastidar, M.G., Mosiolek, M., Bleckmann, A., Dresselhaus, T., Nodine, M.D., and Maizel, A. (2016). Sensitive whole mount in situ localization of small RNAs in plants. The Plant Journal 88: 694–702.

Dello Ioio, R., Linhares, F.S., Scacchi, E., Casamitjana-Martinez, E., Heidstra, R., Costantino, P., and Sabatini, S. (2007). Cytokinins determine Arabidopsis root-meristem size by controlling cell differentiation. Curr Biol 17: 678–682.

Dello Ioio, R., Nakamura, K., Moubayidin, L., Perilli, S., Taniguchi, M., Morita, M.T., Aoyama, T., Costantino, P., and Sabatini, S. (2008). A genetic framework for the control of cell division and differentiation in the root meristem. Science 322: 1380–1384.

Deplancke, B. (2004). A Gateway-Compatible Yeast One-Hybrid System. Genome Res. 14: 2093–2101.

Donner, T.J., Sherr, I., and Scarpella, E. (2009). Regulation of preprocambial cell state acquisition by auxin signaling in Arabidopsis leaves. Development 136: 3235–3246.

Eliasson, L., Bertell, G., and Bolander, E. (1989). Inhibitory Action of Auxin on Root Elongation Not Mediated by Ethylene. Plant Physiol 91: 310–314.

Fahlgren, N., Montgomery, T.A., Howell, M.D., Allen, E., Dvorak, S.K., Alexander, A.L., and Carrington, J.C. (2006). Regulation of AUXIN RESPONSE FACTOR3 by TAS3 ta-siRNA affects developmental timing and patterning in Arabidopsis. Curr. Biol. 16: 939–944.

Garcia, D., Collier, S.A., Byrne, M.E., and Martienssen, R.A. (2006). Specification of leaf polarity in Arabidopsis via the trans-acting siRNA pathway. Curr Biol 16: 933–938.

Gaudinier, A. et al. (2011). Enhanced Y1H assays for Arabidopsis. Nat. Methods 8: 1053–1055.

Gendrel, A.-V., Lippman, Z., Martienssen, R., and Colot, V. (2005). Profiling histone modification patterns in plants using genomic tiling microarrays. Nat. Methods 2: 213–218.

Gutierrez, L., Bussell, J.D., Pacurar, D.I., Schwambach, J., Pacurar, M., and Bellini, C. (2009). Phenotypic Plasticity of Adventitious Rooting in Arabidopsis Is Controlled by Complex Regulation of AUXIN RESPONSE FACTOR Transcripts and MicroRNA Abundance. The Plant Cell 21: 3119–3132.

Hardtke, C.S. and Berleth, T. (1998). The Arabidopsis gene MONOPTEROS encodes a transcription factor mediating embryo axis formation and vascular development. The EMBO Journal 17: 1405–1411.

Heidstra, R. and Sabatini, S. (2014). Plant and animal stem cells: similar yet different. Nature Reviews Molecular Cell Biology 15: 301–312.

Hunter, C., Sun, H., and Poethig, R.S. (2003). The Arabidopsis heterochronic gene ZIPPY is an ARGONAUTE family member. Curr Biol 13: 1734–1739.

Hunter, C., Willmann, M.R., Wu, G., Yoshikawa, M., de la Luz Gutiérrez-Nava, M., and Poethig, S.R. (2006). Trans-acting siRNA-mediated repression of ETTIN and ARF4 regulates heteroblasty in Arabidopsis. Development 133: 2973–2981.

Karimi, M., Bleys, A., Vanderhaeghen, R., and Hilson, P. (2007). Building blocks for plant gene assembly. Plant Physiol 145: 1183–1191.

Kasschau, K.D., Xie, Z., Allen, E., Llave, C., Chapman, E.J., Krizan, K.A., and Carrington, J.C. (2003). P1/HC-Pro, a viral suppressor of RNA silencing, interferes with Arabidopsis development and miRNA unction. Dev. Cell 4: 205–217.

Lampropoulos, A., Sutikovic, Z., Wenzl, C., Maegele, I., Lohmann, J.U., and Forner, J. (2013). GreenGate - A Novel, Versatile, and Efficient Cloning System for Plant Transgenesis. PLOS ONE 8: e83043.

Livak, K.J. and Schmittgen, T.D. (2001). Analysis of relative gene expression data using real-time quantitative PCR and the 2(-Delta Delta C(T)) Method. Methods 25: 402–408.

Mähönen, A.P., ten Tusscher, K., Siligato, R., Smetana, O., Díaz-Triviño, S., Salojärvi, J., Wachsman, G., Prasad, K., Heidstra, R., and Scheres, B. (2014). PLETHORA gradient formation mechanism separates auxin responses. Nature 515: 125–129.

Mallory, A.C. and Vaucheret, H. (2006). Functions of microRNAs and related small RNAs in plants. Nat. Genet. 38 Suppl: S31–36.

Marin, E., Jouannet, V., Herz, A., Lokerse, A.S., Weijers, D., Vaucheret, H., Nussaume, L., Crespi, M.D., and Maizel, A. (2010). miR390, Arabidopsis TAS3 tasiRNAs, and their AUXIN RESPONSE FACTOR targets define an autoregulatory network quantitatively regulating lateral root growth. Plant Cell 22: 1104–1117.

Mecchia, M.A., Debernardi, J.M., Rodriguez, R.E., Schommer, C., and Palatnik, J.F. (2013). MicroRNA miR396 and RDR6 synergistically regulate leaf development. Mechanisms of Development 130: 2–13.

Megraw, M., Baev, V., Rusinov, V., Jensen, S.T., Kalantidis, K., and Hatzigeorgiou, A.G. (2006). MicroRNA promoter element discovery in Arabidopsis. RNA 12: 1612–1619.

Megraw, M. and Hatzigeorgiou, A.G. (2010). MicroRNA Promoter Analysis. Methods Mol. Biol. 592: 149–161.

Okushima, Y. et al. (2005). Functional genomic analysis of the AUXIN RESPONSE FACTOR gene family members in Arabidopsis thaliana: unique and overlapping functions of ARF7 and ARF19. Plant Cell 17: 444–463.

O’Malley, R.C., Huang, S.-S.C., Song, L., Lewsey, M.G., Bartlett, A., Nery, J.R., Galli, M., Gallavotti, A., and Ecker, J.R. (2016). Cistrome and Epicistrome Features Shape the Regulatory DNA Landscape. Cell 166: 1598.

Paque, S. and Weijers, D. (2016). Q&A: Auxin: the plant molecule that influences almost anything. BMC Biology 14: 67.

Plavskin, Y., Nagashima, A., Perroud, P.-F., Hasebe, M., Quatrano, R.S., Atwal, G.S., and Timmermans, M.C.P. (2016). Ancient trans-Acting siRNAs Confer Robustness and Sensitivity onto the Auxin Response. Dev. Cell 36: 276–289.

Poethig, R.S., Peragine, A., Yoshikawa, M., Hunter, C., Willmann, M., and Wu, G. (2006). The function of RNAi in plant development. Cold Spring Harb. Symp. Quant. Biol. 71: 165–170.

Przemeck, G.K.H., Mattsson, J., Hardtke, C.S., Sung, Z.R., and Berleth, T. (1996). Studies on the role of the Arabidopsis gene MONOPTEROS in vascular development and plant cell axialization. Planta 200: 229–237.

Rademacher, E.H., Möller, B., Lokerse, A.S., Llavata Peris, C.I., van den Berg, W., and Weijers, D. (2011). A cellular expression map of the Arabidopsis AUXIN RESPONSE FACTOR gene family. Plant J. 68: 597–606.

Rodriguez, R.E., Ercoli, M.F., Debernardi, J.M., Breakfield, N.W., Mecchia, M.A., Sabatini, M., Cools, T., Veylder, L.D., Benfey, P.N., and Palatnik, J.F. (2015). MicroRNA miR396 Regulates the Switch between Stem Cells and Transit-Amplifying Cells in Arabidopsis Roots. The Plant Cell 27: 3354–3366.

Rodriguez, R.E., Mecchia, M.A., Debernardi, J.M., Schommer, C., Weigel, D., and Palatnik, J.F. (2010). Control of cell proliferation in Arabidopsis thaliana by microRNA miR396. Development 137: 103–112.

Scheres, B. (2007). Stem-cell niches: nursery rhymes across kingdoms. Nature Reviews Molecular Cell Biology 8: 345–354.

Schindelin, J. et al. (2012). Fiji: an open-source platform for biological-image analysis. Nat. Methods 9: 676–682.

Schlereth, A., Möller, B., Liu, W., Kientz, M., Flipse, J., Rademacher, E.H., Schmid, M., Jürgens, G., and Weijers, D. (2010). MONOPTEROS controls embryonic root initiation by regulating a mobile transcription factor. Nature 464: 913–916.

Smet, I.D. et al. (2010). Bimodular auxin response controls organogenesis in Arabidopsis. PNAS 107: 2705–2710.

Stauffer, E. and Maizel, A. (2014). Post-transcriptional regulation in root development. Wiley Interdiscip Rev RNA 5: 679–696.

Sunkar, R., Kapoor, A., and Zhu, J.-K. (2006). Posttranscriptional induction of two Cu/Zn superoxide dismutase genes in Arabidopsis is mediated by downregulation of miR398 and important for oxidative stress tolerance. The Plant Cell 18: 2051–2065.

Thimann, K.V. (1937). On the Nature of Inhibitions Caused by Auxin. American Journal of Botany 24: 407–412.

Tian, C., Muto, H., Higuchi, K., Matamura, T., Tatematsu, K., Koshiba, T., and Yamamoto, K.T. (2004). Disruption and overexpression of auxin response factor 8 gene of Arabidopsis affect hypocotyl elongation and root growth habit, indicating its poss… - PubMed - NCBI. The Plant Journal 40: 333–343.

Truernit, E., Bauby, H., Dubreucq, B., Grandjean, O., Runions, J., Barthelemy, J., and Palauqui, J.C. (2008). High-Resolution Whole-Mount Imaging of Three-Dimensional Tissue Organization and Gene Expression Enables the Study of Phloem Development and Structure in Arabidopsis. The Plant cell 20: 1494–1503.

Wang, J.-W., Czech, B., and Weigel, D. (2009). miR156-regulated SPL transcription factors define an endogenous flowering pathway in Arabidopsis thaliana. Cell 138: 738–749.

Weigel, D. and Glazebrook, J. (2002). Arabidopsis: A Laboratory Manual (CSHL Press).

Weijers, D., Schlereth, A., Ehrismann, J.S., Schwank, G., Kientz, M., and Jürgens, G. (2006). Auxin triggers transient local signaling for cell specification in Arabidopsis embryogenesis. Dev. Cell 10: 265–270.

Wu, G., Park, M.Y., Conway, S.R., Wang, J.-W., Weigel, D., and Poethig, R.S. (2009). The sequential action of miR156 and miR172 regulates developmental timing in Arabidopsis. Cell 138: 750–759.

Yamasaki, H., Hayashi, M., Fukazawa, M., Kobayashi, Y., and Shikanai, T. (2009). SQUAMOSA Promoter Binding Protein-Like7 Is a Central Regulator for Copper Homeostasis in Arabidopsis. The Plant Cell 21: 347–361.

Yoon, E.K., Kim, J.-W., Yang, J.H., Kim, S.-H., Lim, J., and Lee, W.S. (2014). A Molecular framework for the differential responses of primary and lateral roots to auxin in Arabidopsis thaliana. J. Plant Biol. 57: 274–281.

Yoon, E.K., Yang, J.H., Lim, J., Kim, S.H., Kim, S.-K., and Lee, W.S. (2010). Auxin regulation of the microRNA390-dependent transacting small interfering RNA pathway in Arabidopsis lateral root development. Nucleic Acids Res. 38: 1382–1391.

Yu, N., Niu, Q.-W., Ng, K.-H., and Chua, N.-H. (2015). The role of miR156/SPLs modules in Arabidopsis lateral root development. Plant J. 83: 673–685.

